# Zooplankton as a transitional host for *Escherichia coli* in freshwater

**DOI:** 10.1101/2021.12.23.474077

**Authors:** Andrea di Cesare, Francesco Riva, Noemi Colinas, Giulia Borgomaneiro, Sara Borin, Pedro J. Cabello-Yeves, Claudia Canale, Nicholas Cedraro, Barbara Citterio, Elena Crotti, Gianmarco Mangiaterra, Francesca Mapelli, Vincenzo Mondino, Carla Vignaroli, Walter Quaranta, Gianluca Corno, Diego Fontaneto, Ester M Eckert

## Abstract

This study shows that *Escherichia coli* can be temporarily enriched in zooplankton in natural conditions and that these bacteria can belong to different phylogroups and sequence types including environmental as well as clinical and animal isolates. We isolated 10 *E. coli* strains and sequenced the genomes of two of them. Phylogenetically the two isolates were closer to strains isolated from poultry meat than with freshwater *E. coli*, albeit their genomes were smaller than those from poultry. After isolation and fluorescent protein tagging of strains ED1 and ED157 we show that *Daphnia* sp. can take up these strains and release them alive again, thus forming a temporary host for *E. coli*. In a chemostat experiment we show that the association does not prolong the bacterial long-term survival, but that at low abundances it does also not significantly reduce the bacterial numbers. We demonstrate that *E. coli* does not belong to the core microbiota of *Daphnia*, suffers from competition by the natural microbiota of *Daphnia*, but can profit from its carapax to survive in water. All in all, this study suggests that the association of *E. coli* to *Daphnia* is only temporary but that the cells are viable therein and this might allow encounters with other bacteria for genetic exchange and potential genomic adaptations to the freshwater environment.

**Importance:** The contamination of freshwaters with faecal derived bacteria is of major concern regarding drinking water acquisition and recreational activities. Ecological interactions promoting their persistence are still very scarcely studied. This study, which analyses the survival of *E. coli* in the presence of zooplankton, is thus of ecological as well as water safety relevance.

## 1. Introduction

Faecal bacteria can enter aquatic environments by different routes, e.g., sewage discharge or direct faecal deposition (Korajkic, et al. 2019). Although faecal bacteria are tendentially seen to rapidly drop in abundance once outside host, some aquatic environments might allow their long-term survival or growth (Korajkic, et al. 2019). *Escherichia coli* is one of the predominant species forming the microbial community that shapes the commensal gut of vertebrates (Martinson and Walk 2020). Thus it is commonly released, by faecal route, into aquatic environments (Espinosa-Urgel and Kolter 1998), and it is therefore used as a faecal indicator bacterium (FIB) to evaluate anthropogenic water pollution (Jang, et al. 2017). Clinically relevant *E. coli* strains (Vignaroli, et al. 2013; Jørgensen, et al. 2017), including antibiotic resistant isolates (Vignaroli, et al. 2012; Baniga, et al. 2020; Fernandes, et al. 2020), can be found in waters. Moreover, there is evidence that *E. coli* can adapt to a freshwater lifestyle shown for example through differential gene expression once incubated in water (Espinosa-Urgel and Kolter 1998).

Naturalized *E. coli*, e.g. *E. coli* that entered the aquatic environment from the gut and then adapted to this lifestyle, were isolated from lake sediments and phytoplankton and different studies showed the capability of this species to genetically adapt and persist in the environment for example through encapsulation (Power, et al. 2005; Ishii, Ksoll, et al. 2006; Ishii, Yan, et al. 2006; Walk, et al. 2007). Several *E. coli* isolated from freshwaters had a small genome size and other peculiarities at the genomic level that suggested an evolutionary adaptation to this habitat (Touchon, et al. 2020). This is particularly interesting since genome reduction has been proposed repeatedly as an adaptation to aquatic environments in common environmental bacteria (Grote, et al. 2012; Salcher, et al. 2019) and in experimental systems (Lee and Marx 2012; Baumgartner, et al. 2017). Thus, the aquatic environment may contribute to the genetic evolution of *E. coli* (Touchon, et al. 2020).

However, mammal associated faecal bacteria usually persist badly in cold habitats, such as deep lakes. Particularly, pelagic cold waters are a very hostile environment for such gut symbionts. If they are not grazed by flagellated predators upon arrival, they are surely not favoured by resource competition (González, et al. 1992; Wanjugi and Harwood 2013; Eckert, et al. 2019). Since evolution, as seen in the genome reduction of *E. coli* (Touchnon, et al. 2020), takes time, a certain long-term persistence of vertebrate commensal strains in the aquatic habitat is crucial, and the question remains: In which niche does persistence take place? In clinical settings these bacteria thrive better in biofilms (Costerton, et al. 1999) and they might persist in a similar niche in aquatic habitats (Hall-Stoodley and Stoodley 2005; Eckert, et al. 2018). In lake environment biofilms can be formed on dead organic and inorganic material, sediments and stones or animals, and FIB have been found in sediments (Haller, et al. 2009; John, et al. 2009), macrophytes (Quero, et al. 2015) and on fish (Declerck, et al. 2007; Abgottspon, et al. 2014) for example. Much less attention has been devoted to small invertebrates, i.e. zooplankton, as a potential host for such bacteria. Such animals are interesting since their microbiota seems to be composed by many transient microbes and thus likely more prone to be invaded by allochthonous bacteria (Grossart, et al. 2010; Hammer, et al. 2019; Callens, et al. 2020; Eckert, et al. 2021). In fact, antibiotic resistant bacteria were easily removed from the surrounding water in a laboratory experiment but persisted in the crustacean *Daphnia obtusa* (Eckert, et al. 2016) and FIB have even been shown to even exchange genetic material in *Daphnia pulex* (Olanrewaju, et al. 2019). It is generally assumed that the presence of *Daphnia* sp. reduces *E. coli* abundance in the water (Burnet, et al. 2017; Ismail, et al. 2019). Nevertheless, in a study based on 16S rRNA gene amplicon sequencing from a lake *E. coli/Shigella* made up for a large percentage of the copepod and *Daphnia* microbiota, but was present only with low abundances on stones, water and sediments (Eckert, et al. 2020). In the present study we thus wanted to investigate the nature of the relationship between *E. coli* and *Daphnia* in the freshwater environment to clarify the possible role that *Daphnia* might have on the persistence of *E. coli* in waters from an ecological point of view.

Here we tested the hypothesis that an association with zooplankton animals of the genus *Daphnia* could help a FIB, *E. coli*, to survive in the harsh conditions of a freshwater lake. It was shown that the presence of *Daphnia pulex* in a few hours reduced the abundance of surrounding *E. coli* (Burnet, et al. 2017) but here we were interested in the longer-term association of the bacteria with the animal under natural conditions. Our hypothesis is that such association might help the bacterium to adapt to this environment, leading to evolution in its genome. Therefore, we quantified *uid*A, an indicator gene of *E. coli*, in DNA extracted from various potential niches for FIB from a freshwater lake, including stone biofilms, zooplankton, sediment and compared it to the pelagic water. Moreover, we searched for the occurrence of *E. coli* related 16S rRNA reads in a large dataset of zooplankton associated microbiota. We then isolated *E. coli* strains from a *Daphnia* host, genotyped them and tagged two of the strains with fluorescent proteins and sequenced their genome. This allowed us to conduct experiments on the association of these strains with an invertebrate host. Our hypothesis was that despite *Daphnia* grazing would reduce the abundance of *E. coli* in the water it would still allow for a better survival of the FIB over a longer time thanks to a short-term refuge of part of the population within its gut. Moreover, we speculate that such an association might in a long term help *E. coli* to adapt to freshwater over physiological and/or genetic adaptations.

## 2. Methods

### 2.1 E. coli *abundance in zooplankton microbiomes*

Initially, we observed a large number of *E. coli/Shigella* related reads in our Illumina MiSeq dataset of zooplankton (*Daphnia* gr. *galeata/longispina* and copepod) associated microbiota obtained from Lake Maggiore (Eckert, et al. 2020). In order to confirm the presence of *E. coli*, quantitative PCR (qPCR) assays were conducted using *E. coli* specific primers for the *uid*A gene (1-CAATGGTGATGTCAGCGTT and 2-ACACTCTGTCCGGCTTTTG, (Srinivasan, et al. 2011)) using the RT-thermocycler CFX Connect (Bio-Rad). The standard calibration curve for the quantification of *uid*A was carried out as described elsewhere (Sabatino, et al. 2015) and the gene concentration was expressed as gene *copy/Daphnia* or mL of water. The same was done for *Daphnia obtusa* isolated from a small pond in the garden of CNR-IRSA (Eckert, et al. 2016). Twenty individuals were washed in autoclaved *MilliQ* water, recollected per triplicate and introduced in DNA Isolation Kit in Ultra Clean Microbial or Power Soil DNA Isolation Kit (Qiagen) for DNA extraction. The right size of all qPCR products was evaluated by electrophoresis (30 min at 80 V, 1% agarose gel). The efficiency of reaction was 87.5% and R^2^ was 0.99. The limit of quantification (LOQ) was determined (Bustin, et al. 2009) and was 45 gene copy/μL.

Furthermore, we checked a large dataset of microbial communities associated with zooplankton, from many natural freshwater habitats and cultures published elsewhere (Eckert, et al. 2021) looking for the presence of *E. coli/Shigella* affiliated reads. The dataset is composed of cladocerans (*Daphnia magna, Daphnia obtusa, Diaphanosoma brachyurum, Simocephalus* sp. and *Mesocyclops leukarti)* and rotifers (*Adineta vaga, Keratella serrulata, Lecane elsa, Lecane inermis, Epiphanes senta, Keratella quadrata* and *Polyarthra* sp.) (see also Figure 2).

### 2.2 E. coli *isolates*

#### 2.2.1 Isolation

Individuals of *Daphnia obtusa* were collected two or three times per week between May to July and October to November, from a rainwater-fed pond in the garden of the CNR-IRSA in Verbania (Italy). Thirty individuals of *Daphnia obtusa* were washed in autoclaved *MilliQ* water (Millipore), crushed and sonicated (3 times, 1 minute each cycle with vortex within cycles) in 1 ml of physiological solution. Serial ten-fold dilutions were performed (from 1:10 to 1:10^6^). 1 ml of each dilution was filtered onto nitrocellulose membrane filters (type GSWP, 25 mm diameter, 0.22 μm pore size, Millipore) and filters were plated onto mFC agar plates (Biolife) and incubated for 48 h at 37 ºC.

#### 2.2.2 *Identification and genetic characterization of* E. coli *isolates*

Aliquots of presumptive *E. coli* colonies were introduced in 1 mL of physiological solution, centrifuged (5000 rcf, 4ºC, for 10 minutes), boiled for 15 minutes, frozen for 2-4 hours and centrifuged as before. DNA from presumptive *E. coli* colonies and from *Daphnia* were tested for the presence of the *uid*A gene using the primers above mentioned by PCR. Conditions for the PCR assays were: 5 μl of Buffer 5X, 0.5 μl dNTPs (10 mM), 0.2 μl Taq-polimerase (5U/ μl), 0.25 μl per each primers (100 μM), and water that it was added to arrive to final volume of 23 μl. PCR reactions were: denaturation 3 min at 95ºC, 35 cycles of 30 s at 95ºC, 1 min at 58ºC and 1 min at 72ºC, and a final extension step of 7 min at 72ºC. PCR products were separated by agarose gel electrophoresis (1%) and visualized with GelRed (Midore Green Advance DNA Stain). In order to assign a specific phylogroup or clade to the 10 *E. coli* strains isolated we used the PCR-based method described by Clermont et al., 2013 (Clermont, 2013). PCR products were separated by agarose gel electrophoresis (1%) and visualized with GelRed (Midori Green Advance DNA Stain). The 10 *E. coli* isolates were further analyzed to obtain an unambiguous DNA fingerprint by Enterobacterial Repetitive Intergenic consensus (ERIC-PCR) as previously described (Versalovic, et al. 1991). ERIC-PCR products were separated by electrophoresis for 8 h at 40 V/cm, in 2% agarose Tris borate-EDTA (TBE) gel stained with GelRed (Midori Green Advance DNA Stain).

The strains ED1, ED4, ED8, ED157 and ED166 were chosen to be typed by Multi-Locus Sequence Typing (MLST) by sequence analysis of internal fragments of seven housekeeping genes (adk, fumC, gyrB, icd, mdh, purA, recA) according to the Achtman scheme (http://enterobase.warwick.ac.uk/species/index/ecoli). The allelic profiles of the seven genes and the resulting Sequence Type (ST) were determined from the submission of sequence data on the PubMLST database (https://pubmlst.org).

#### 2.2.3 Pathogenicity assay

The hemolytic activity of the strains was evaluated as described by Ghosh et al. (2014) with some modifications. Briefly, 4 mL of freshly drawn, heparinized human blood was diluted with 25 mL of phosphate buffered saline (PBS), pH 7.4. After washing three times in 25 mL of PBS, the pellet was resuspended in PBS to 20 vol %. A 100 μL amount of erythrocyte suspension was added to 100 μL of bacterial strains. PBS and 0.2% Triton X-100 were used as the negative and positive controls, respectively. After 1 h of incubation at 37 °C each well was centrifuged at 1200 × g for 15 min, the supernatant was diluted 1:3 in PBS and transferred to a new plate. The OD350 was determined using the Synergy HT microplate reader spectrophotometer (BioTek, Winooski, VT, USA). The hemolysis (%) was determined as follows:

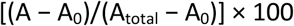

where A is the absorbance of the test well, A_0_ the absorbance of the negative control, and A_total_ the absorbance of the positive control; the mean value of three replicates was recorded.

To detect the biofilm development, the strains were grown in LB medium (Oxoid), adjusted to 5 × 10^6^ CFU/mL and inoculated (100 μL) in 24-well polystyrene plates (VWR). After 24 h of incubation at 37 °C, and 24°C the wells were washed with PBS to eliminate unattached cells and covered with 0.1% (v/v) crystal violet (CV) dissolved in H_2_O for 15 min, washed in PBS and air-dried. The remaining CV was dissolved in 85%ethanol for 15 min at room temperature and 200 μL from each well was transferred to a 24-well plate for spectrophotometric quantification at 570 nm (Multiscan Ex Microplate Reader, Thermo Scientific, Waltham, MA, USA. The strains were classified as strong, moderate, or weak biofilm producers based upon the ODs of the bacterial biofilms (Stepanović, et al. 2007). Quantification of biofilm in microtiter plates: overview of testing conditions and practical recommendations for assessment of biofilm production by staphylococci (Stepanović, et al. 2007). All assays were performed in triplicate using independent cultures.

#### 2.2.3 Genome sequencing and analysis

Two strains, namely strains ED1 and ED157, were chosen for genome sequencing. These two strains were selected because they were both affiliated with the D phylotype and because of their respective sequence type: ED1 belonged to a sequence type that contained many bacteria isolated from mammals including humans. The sequence type of ED157 only contained another *E. coli* strain that was isolated from water. The strains were grown in Luria Broth (LB, Merck Life Science) overnight and DNA extraction was performed using the ultraclean microbial DNA extraction kit (QIAGEN).

Purified DNA was sequenced on a NovaSeq Illumina Platform (IGA technologies, Padova, Italy), providing a total of 10 and 15 million reads of output for ED1 and ED157 strains, respectively. One of the two genomes was already mentioned in a previous article (Riva, et al. 2020). Briefly, reads were first trimmed using Trimmomatic (Bolger, et al. 2014) and genomes were assembled using SPAdes default parameters (Prjibelski, et al. 2020), obtaining a total of 54 and 59 assembled contigs >1 Kb, respectively.

To verify the phylotype of *E. coli* strains ED1 and ED157, we submitted the genome sequences to the website ClermoTyper (Beghain, et al. 2018). Plasmids’ presence in the genomes was examined through the platform PlasmidFinder (Carattoli, et al. 2014), while the identification of antimicrobial resistance genes was done using ResFinder4.0 (Bortolaia, et al. 2020). Virulence genes were instead found using the VirulenceFinder 2.0 platform (Joensen, et al. 2014) and phage genome sequences were recognized using PHASTER (Arndt, et al. 2016).

In order to evaluate whether the *Daphnia* associated isolates were similar to other freshwater *E. coli* we compared them to other genomes from the D phylotype mentioned in Touchon et al. (2020): i) C4_38 and C2_45 strains, isolated from poultry meat, and ii) E5895 and E6003 strains, isolated from freshwater (Touchon et al., 2020; Tab. 1). Strains E5895 and E6003 were indeed randomly selected as representative *E. coli* strains adapted to the freshwater environment, owing a reduced genome, while the other two strains (C4_38 and C2_45) were randomly selected as representatives of strains from poultry meat, which are known to have the largest average genome within the *E. coli* species (Touchon et al., 2020).

**Tab. 1.**
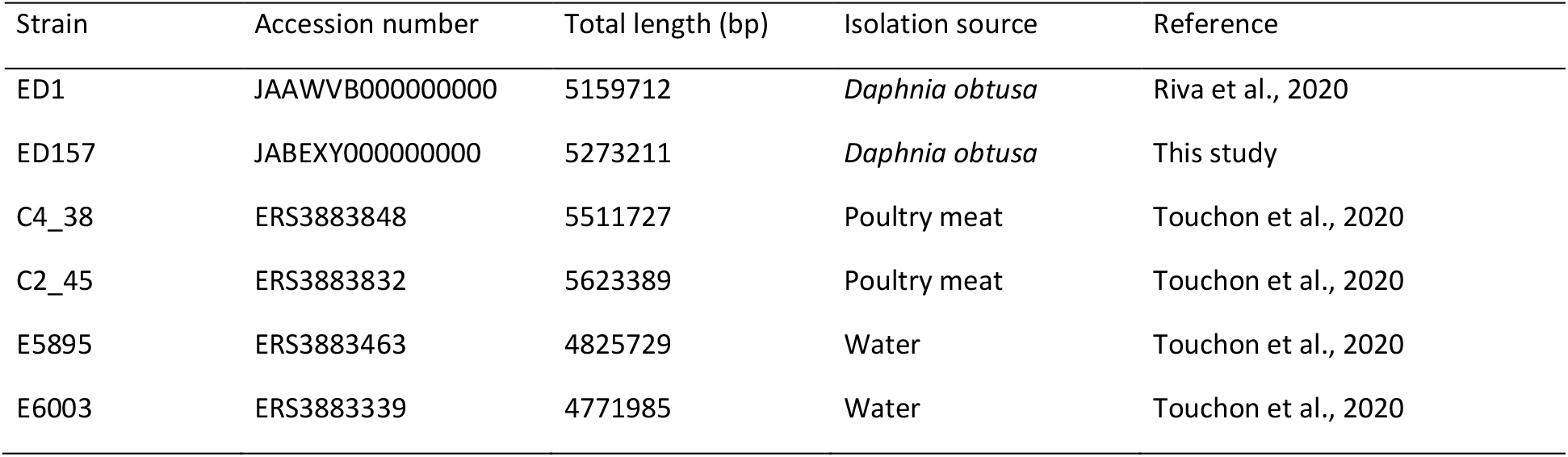
*Escherichia coli* genomes included in the study, their length, the isolation source, and the reference where they were first reported.

| Strain | Accession number | Total length (bp) | Isolation source | Reference |
| --- | --- | --- | --- | --- |
| ED1 | JAAWVB000000000 | 5159712 | *Daphnia obtusa* | Riva et al., 2020 |
| ED157 | JABEXY000000000 | 5273211 | *Daphnia obtusa* | This study |
| C4_38 | ERS3883848 | 5511727 | Poultry meat | Touchon et al., 2020 |
| C2_45 | ERS3883832 | 5623389 | Poultry meat | Touchon et al., 2020 |
| E5895 | ERS3883463 | 4825729 | Water | Touchon et al., 2020 |
| E6003 | ERS3883339 | 4771985 | Water | Touchon et al., 2020 |

Phylogenetic analysis considering the whole genome sequences was performed through the MICROSCOPE platform (http://www.genoscope.cns.fr/agc/microscope (Blondel, et al. 2008; Vallenet, et al. 2009)). The phylogenetic tree was built with “genome clustering” MICROSCOPE tool. Genomic similarity was estimated using Mash, with distances correlated to ANI like: D ≈ 1-ANI. From all the pairwise distances of the genomes set, a tree was constructed dynamically using the neighbor-joining javascript package, displaying clustering annotations. This clustering was computed from all-pairs distances ≤ 0.06 (≈94% ANI), which correspond to the ANI standard to define a species group. Clustering was computed using the Louvain Community Detection.

In order to evidence differences and examine the distribution of protein families across the *E. coli* genomes indicated in Tab. 1, we used the “Protein Family Sorter” tool of PATRIC (https://www.patricbrc.org/), setting Genus-specific families (PLfams) (Davis, et al. 2016; Davis, et al. 2020).

#### 2.2.5 GFP and DsRed tagging of strains

Competent cells of *E. coli* ED1 and ED157 strains were prepared in LB medium following the protocol described in Favia et al. (2007). Sixty microliters of competent cells (~10^10^ cells ml^-1^) were mixed with 100-200 ng of plasmid DNA, transferred to a cold 0.1-cm-diameter cuvette and pulsed at 1,700 V in the Electroporator 2510 apparatus (Eppendorf, Milan, Italy). Plasmids used were pHM2-Gfp (Favia, 2007) and pKan(DsRed) (Crotti, et al. 2009). Following the pulse, cells were immediately supplemented with 1 ml of LB medium and incubated at 37°C for 1 h. Transformants were selected by plating on LB agar medium added with i) 100 μg·ml^-1^ kanamycin (KMY), 40 μg·ml^-1^ 5-bromo-4-chloro-3-indolyl-b-D-galactopyranoside (X-Gal) and 0.5 mM isopropyl-b-D-thiogalactopyranoside (IPTG) in case of plasmid pHM2-Gfp or ii) 100 μg·ml^-1^ KMY in case of plasmid pKan(DsRed). Verifications of the presence of pHM2-Gfp or pKan(DsRed) plasmids were done observing the bacterial cells by fluorescence microscopy. Furthermore, the identity of *E. coli* transformants was confirmed by BOX-PCR amplification (Urzì, et al. 2001) by comparing the BOX-PCR profiles with those of wild type ED1 and ED157 strains.

### 2.4 *Daphnia-E. coli* association experiments

All laboratory experiments were carried out using *Daphnia obtusa* from the garden of CNR-IRSA. The animals were collected 2 days before the experiment to adapt them to lab conditions. They were washed with artificial lake water medium (ALW, inorganic compounds in composition described here: (Zotina, et al. 2003)) and fed with a small amount of washed *Kirchneriella* sp. and kept in the dark before experimental use. The animals were washed again in ALW before each experiment and the experiments were conducted in the same medium if not specified differently. The *E. coli* strains ED1 and ED157 tagged with the fluorescent proteins which were used in the experiments were grown overnight at 37°C in liquid LB containing 100 μg·ml^-1^ KMYto maintain the plasmid. The strains were centrifuged and washed twice with ALW before inoculation in experimental treatments the next day. All experiments were carried out at room temperature in the dark. Figure 1 illustrates the approaches taken for the presented laboratory experiments. All figures of the experiments were drawn in *R (RCore Team 2013*), using the packages *ggplot*2 (Wickham 2009), *reshape*2 (Wickham 2012) and *cowplot (Wilke 2020)*.

**Figure 1.**
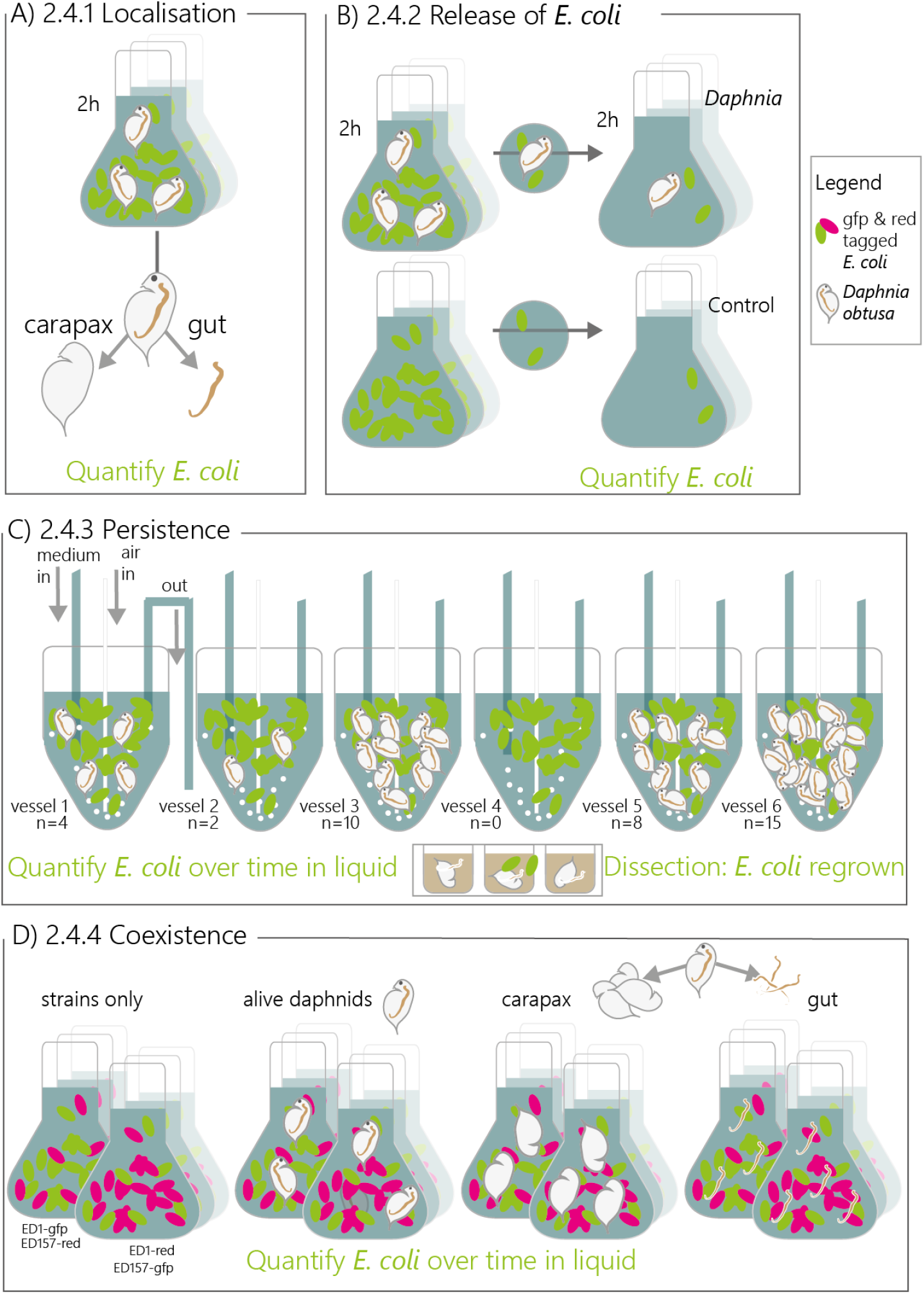
Experimental set-up of all experiments involving *E. coli* and *Daphnia obtusa*. A-D schematically depict the set-up of each experiment with the numbers corresponding to the subtitle within the material and method section 2.4. Experiments in A, B and D were conducted in batches, whereas experiment C was conducted in a chemostat.

#### 2.4.1. Localisation of *E. coli* on *Daphnia*

We verified where *E. coli* attached to *Daphnia* by incubation of *E. coli* with live individuals *Daphnia* for 4 hours. Animals were then dissected and different body parts were subjected to qPCR amplification of the *gfp*-gene to verify the presence and quantify *E. coli*. qPCR assay was carried out in a volume of 20 μL containing 2 μL of DNA, 0.5 μM of each primer (1-GAAGATGGAAGCGTTCAA and 2-AGGTAATGGTTGTCTGGTA, (Hale, et al. 2015), 10 μL of SsoAdvanced universal SYBR Green supermix (Bio-Rad), and filtered and autoclaved MilliQ water (Millipore) to the final volume. The program of qPCR was 95°C for 2 min, 35 cycles of 95 °C for 15 s, 54°C for 30 s and 72 °C for 15 s. Melting curve was performed from 60 °C to 95 °C with increments of 0.5 °C/5 s. The right size of all qPCR amplicons was evaluated by electrophoresis run (carried out as described above for *uid*A gene). The standard curve for the *gfp-gene* quantification was carried out by the dilution of the purified and quantified amplicon, as made for the *uid*A gene and previously described in Sabatino et al. (2015). The efficiency of reaction was 91% and R2 was 0.99. The LOQ (determined as described above for the *uid*A gene) was 9.85 gene copy/μL. The concentration of the *gfp*-gene was expressed as gene copy/daphnia.

#### 2.4.2 Release of *E. coli* after gut passage

We then tested whether *E. coli* was a food source for *Daphnia*, if *Daphnia* functioned as a refuge for *E. coli* or if they simply passed through the gut. First we incubated ED1-gfp and ED157-gfp separately with and without *Daphnia* in 50 ml ALW for 2h in the dark at room temperature, then we transferred 50 μl of only water or water with a single *Daphnia* (+D treatment or control) and, after another 2h of incubation, we compared the amount of ED1-gfp transferred in the surrounding water. ED1-gfp was counted on a flow cytometer as green fluorescent events (BD C6, Accuri). Differences between treatments were evaluated using a linear model on log-transformed count data for the response variable, conducted in *R*.

#### 2.4.3 Persistence of *E. coli* with *Daphnia*

The chemostat was a continuous culturing system with three medium tanks containing ALW attached to six vessels containing 700 ml of medium. A non-axenic *Kirchneriella* culture was added to both medium and vessels at a density of around 20,000 cells per ml (T-5). The system was kept in the dark. After three days (T-2) the chemostat pumps were switched on with a daily water replacement rate of 10%. After one day (T-1) ED1-gfp was added to the vessel at a concentration of 10^6^ cells ml^-1^ as well as algae to maintain around 20,000 cells ml^-1^. This experiment was conducted with strain ED1 because according to its sequence type it is a more relevant potential contamination from mammalian origin into freshwaters. After another day (T0) *Daphnia obtusa* was added to the vessels, which were randomly assigned with a quantity of animals in a gradient with the following number of animals per vessel: 0, 2, 6, 8, 10 and 15. Samples of 40 ml were taken every 2-3 days over the outflow of the chemostat vessel and *Kirchneriella* solution was always added after sampling to maintain food for *Daphnia*. These samples were used for CFU counts for ED1-gfp and microscopy counts for both algae and ED1. For phytoplankton counts 10 ml solution was filtered on 0.45 μm pore-size polycarbonate filters and at least 10 fields and 500 cells were counted at a magnification of 80’000x at an epifluorescence microscope (Zeiss). For CFUs of day 0 and day 2 spots of 5ul of a gradual dilution between 1 and 10^-4^ were spotted on LB+KMY plates and grown for 24h at 37°C and counted. The presence of the gfp and thus univocal identification of ED1-gfp was done by placing the plate on a transilluminator (UV light) and observation of green fluorescence of the colonies. Due to the strong reduction of *E. coli* numbers of CFU for T 5 and 7 were counted by filtration of 1ml of undiluted and 1:10 diluted sample and on T 9 by filtration of 1ml and 10ml of sample, on a 0.2μm pore-size nitrocellulose membrane filter that was placed on the plates and colonies were counted as described above. All spots and filters were done in triplicate per sample. On T12 *Daphnia* numbers had strongly reduced (see Supplementary Table 1) thus the experiment was considered finished. Individuals of *Daphnia* were extracted from the vessels, washed 3 times with sterile ALW and then dissected; the different body parts were placed in 200ul LB+KMY in a black multiwell-plate in a plate reader (GlowMax, Promega). Growth of *E. coli* was detected by monitoring fluorescence over 48 h every 30 min. The total number of adult dissected *Daphnia* was 12 (3 from vessel 6, 4 from vessel 5, 4 from vessel 3 and 1 from vessel 1) and the number of juvenile *Daphnia* was 9 (1 from vessel 6, 4 from vessel 5, 2 from vessel 3); 7 negative controls were included. We checked the influence of the original gradient of abundance of *Daphnia* on the CFU of *E. coli* by a generalised linear model assuming a negative binomial distribution of the data. The model was evaluated using *check_model* from the package *performance* (Lüdecke, et al. 2019) and the output depicted as a type II analysis of variance table using the *car* package (Fox, et al. 2012).

#### 2.4.4 Coexistence experiment

In order to test whether ED1 and ED157 reacted similarly to the presence of *Daphnia* and its associated bacteria we conducted a batch experiment where we incubated both strains together with no *Daphnia*, alive *Daphnia*, and dissected *Daphnia* for which we made one treatment containing the *Daphnia* guts and one containing the *Daphnia* carapax and filtration apparatus and their associated bacteria. Each replicate was amended with either 3 alive *Daphnia* or dissection pieces from 10 *Daphnia* in 2ml in 1:100 diluted LB with ALW and in triplicate. Moreover each treatment was conducted twice once using ED1-*gfp* + ED157-DsR*ed* and once using ED1-DsR*ed* and ED157-*gfp* to account for potential differences in fitness reduction by the two different fluorescence markers (total treatment n= 2 stainings x 3 replicates x 4 treatments = 24). In fact in both cases ED1-DsR*ed* had a fitness advantage, thus the numbers presented here are averages between the CFUs counted for gfp and red of the same strain in the same treatment. CFUs were counted over 10 days starting from T4 on by spotting of 5ul diluted up to 10^-8^ in triplicates and green and red colonies were counted on a trans-illuminator (UV). The experiment was stopped after 10 days due to major mortality of *Daphnia* in the alive treatment (>90%) and data for the first 8 days was plotted. To evaluate the long-term differences between treatments data from March 24 and 26 were used (6 and 8 days). A generalised linear model assuming a negative binomial distribution of the data was made to evaluate the effect of the treatment and the date on the abundance of both *E. coli* ED1 and ED157 (*glm.nb* in R with model: CFU ~ treatment (4 levels: alive, carapax, gut, no *Daphnia*) + strain (2 levels: ED1 or ED157) + date (2 levels: March 24 and 26)). Model check and model output were performed as for the analysis of persistence of *E. coli* with *Daphnia*; in addition, pairwise differences between treatments were evaluated with a post-hoc using *emmeans* from the homonymous package (Lenth 2020).

### 2.5 Data and code accessibility

All scripts and raw data are deposited at https://github.com/EsterME/E_coli_Daphnia. ED1 genome was deposited into the NCBI-Genbank database under accession number JAAWVB00000000 (Riva, et al. 2020). ED 157 Genome Shotgun project has been deposited at DDBJ/ENA/GenBank under the accession JABEXY000000000. The version described in this paper is version JABEXY010000000.

## 3. Results

### 3.1 *Abundance of* E. coli *in Lake Maggiore*

By screening for the presence/abundance of the *E. coli* specific marker gene *uid*A in DNA extracted from three different locations in Lake Maggiore, we found that the gene was absent in the sediments, epilithic biofilms and water samples but could be found in both *Daphnia gr. galeata/longispina* and copepods, showing between 144-976 (mean 580) copies per animal (Figure 2A).

**Figure 2.**
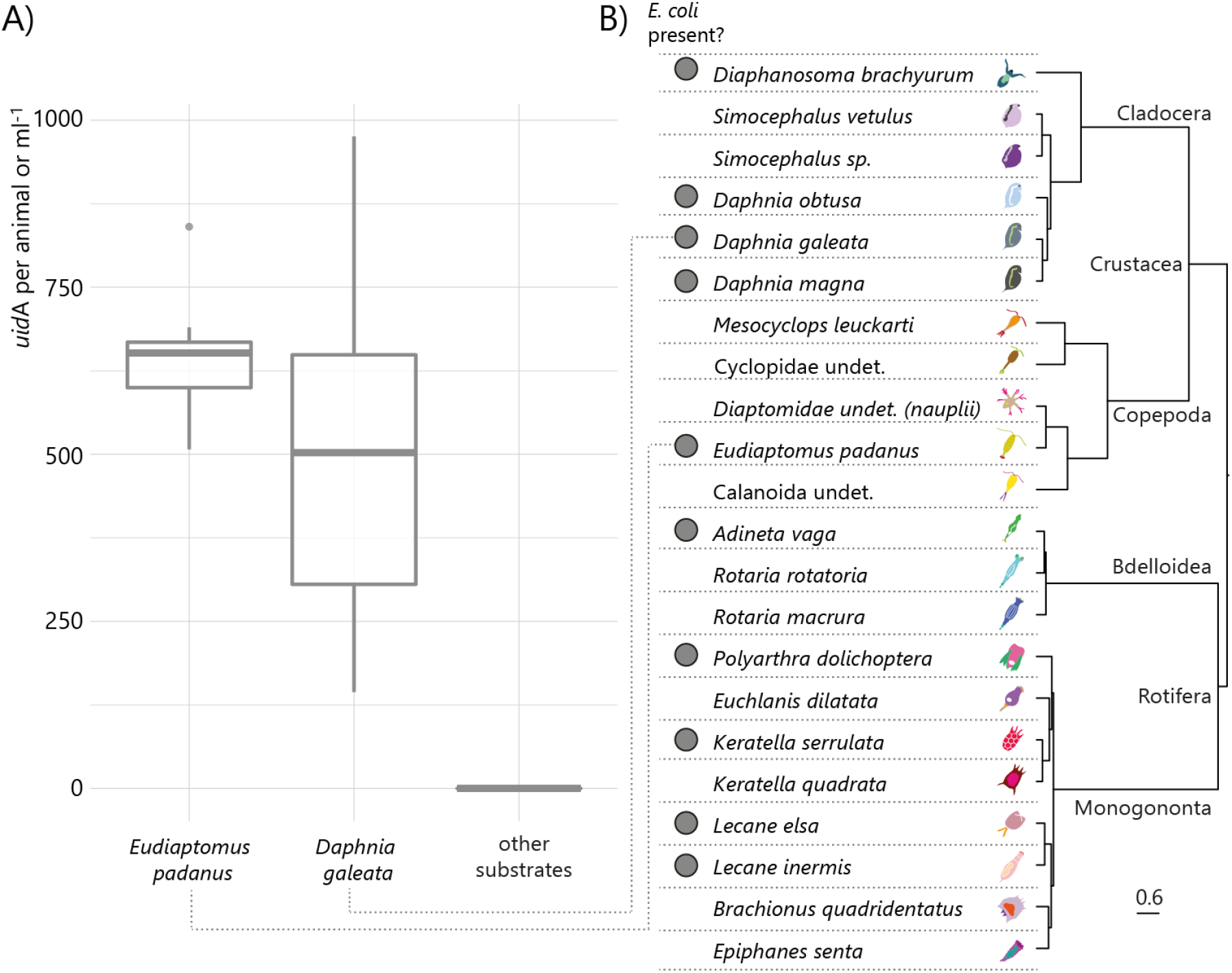
A) Boxplots of the abundance of the *E. coli* specific uid A gene copies in DNA isolated from animals and other substrates that include sediments, stones and water from 10 and 40m depth from Lake Maggiore. For each plot, the tick horizontal line represents the median value, the box includes 50% of the data from the first to the third quartile, the whiskers extend to the minimum and maximum data within the 1.5 interquartile range and the dots represent single outlier data points outside such range. B) Occurrence of *E. coli* in various zooplankton species, a grey dot means *E. coli* was found in the microbiome of at least one sample. Images and phylogeny of animals are given as reference and are modified from Eckert et al, 2021.

### 3.2 E. coli *in other zooplankton microbiomes*

We screened a large dataset of zooplankton-related microbiomes and could find the presence of *E. coli/Shigella* related 16S rRNA gene sequences in samples from other cladocerans (*Daphnia magna, Daphnia obtusa, Diaphanosoma brachyurum)* and rotifers (*Adineta vaga, Keratella serrulata, Lecane elsa, Lecane inermis* and *Polyarthra* sp.) (Figure 2B). *E. coli/Shigella* was not found in other rotifers (*Epiphanes senta, Keratella quadrata), Mesocyclops leukarti*, a large calanoid copepod, and the cladoceran *Simocephalus* sp. (Figure 2B). We quantified *uid*A in *Daphnia obtusa* sampled from a rainfed pond, because their *E. coli/Shigella* related reads were particularly high: we also confirmed the presence of *E. coli uidA* gene by qualitative Real Time PCR.

### 3.3 E. coli *isolates*

#### *Isolation of* E. coli *from* Daphnia obtusa

We attempted to isolate *E. coli* from *Daphnia obtusa* to further investigate which phylogroups of *E. coli* were affiliated with zooplankton. Through multiple isolation campaigns we retrieved ten *E. coli* strains and identified their phylogroup: strains ED1, ED2, ED3, ED4, and ED5 formed one cluster and were affiliated with phylogroup D/E and strains ED157, ED158 and ED166 a second cluster affiliated with phylogroup D/E (we did not succeed in the discrimination between these two phylogroups for these strains), strains ED8 and ED12 to phylogroup B1 (Figure 3A). Five of these strains were further chosen for MLST (ED1, ED4, ED8, ED157 and ED166) and pathogenicity assays: none of the strains showed traits of pathogenicity except for weak biofilm formation for the isolates ED1, ED4 and ED166 and they were classified in four different sequence types (ST38, ST1727, ST3573 and ST4166) (Figure 3A and Supplementary Table 2). We then analysed the strains deposited in the MLST database affiliated with these STs and most of the ST38 isolates were of human origin, whereas ST1727 included strains mostly isolated from animals (41%) than from humans (10%). The other two STs (ST3573 and ST4166) have been rarely described and they included isolates from non-human sources (Figure 3B).

**Figure 3.**
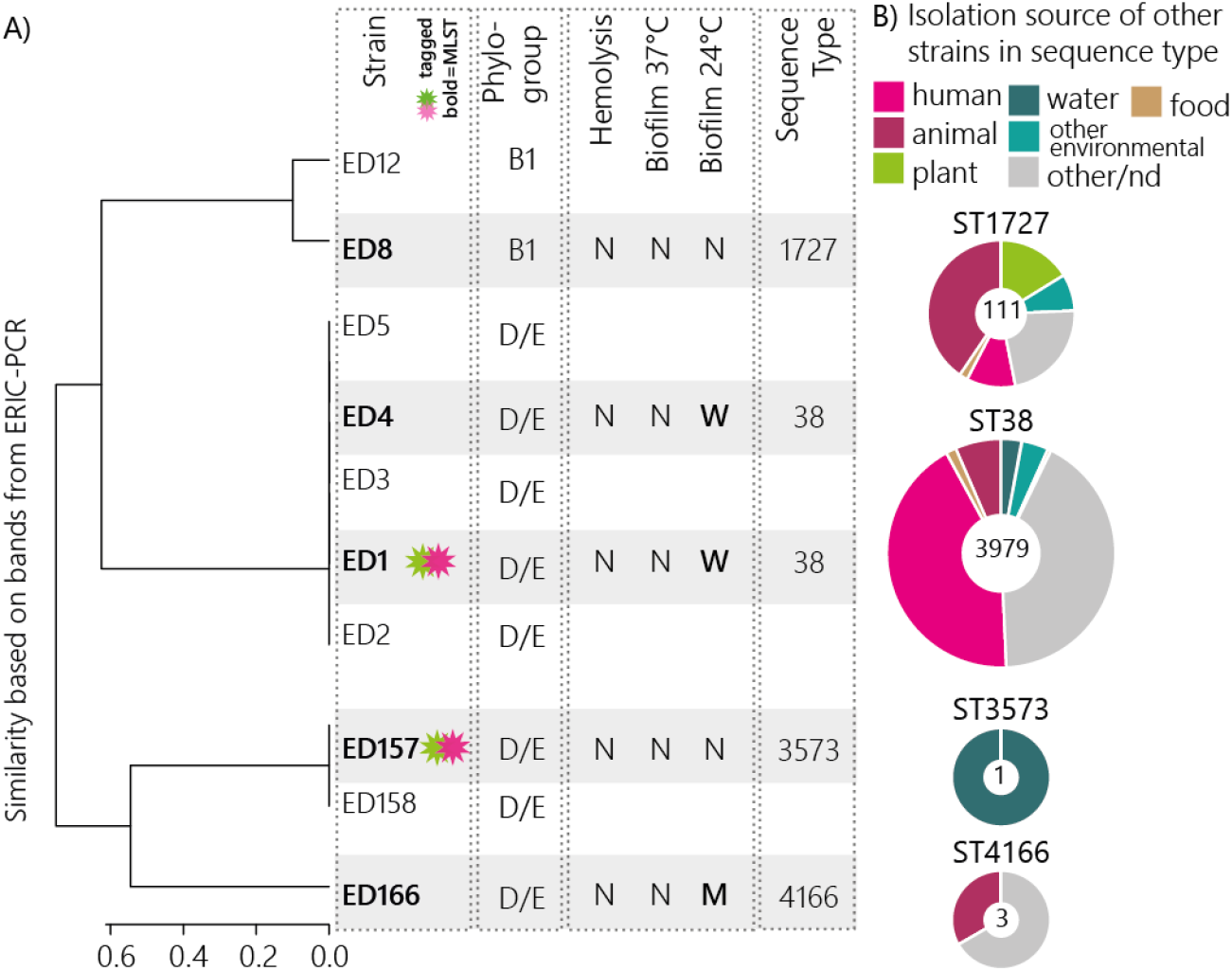
**A.** (fLTR) Cluster dendrogram of dissimilarity of the ERIC profile of the different *E. coli* strains isolated from *D. obtusa* and their phylogroup according to the ERIC profile. Strains that are grey shaded also present data for their phenotype in the pathogenicity assay (N=none, M=medium and W=weak), their sequence type according to Multi-locus sequence typing. **B**. Pie-charts summarising the isolation source of the deposited *E. coli* strains of the same sequence type as the *E. coli* strains from daphnids; the number in the middle of the pie-charts denotes the number of strains deposited per each sequence type.

ED1 and ED157 were selected for further analysis. The rationale behind the selection of these two isolates was their ST: ED1 was affiliated with ST38 where many other *E. coli* seem to be associated with mammals or even pathogens, whereas ED157 was affiliated with ST3573, including only one *E. coli* strain isolated from water. Genomes of ED1 and ED157 strains were sequenced and the strains were successfully marked with gfp and DsRed proteins, for interaction studies.

#### Genome analysis of ED1 and ED157

Genome sequencing and analysis performed by ClermoTyper and phylogenetic tree construction with MICROSCOPE showed that ED1 and ED157 genomes belong to the D phylotype (Figure 4). Phylogenetic analysis performed on the whole genome sequences of the strains and genomes of water and poultry isolated *E. coli* showed that *E. coli* strains isolated from *Daphnia* sp. did not cluster with the water isolates. Conversely, strains isolated from poultry meat clustered with the genomes of the *Daphnia* isolates, whereas the water-related *E. coli* clustered in a sister group (Figure 4). Comparing genome size, we could observe that ED1 and ED157 genome size were bigger than the genome sequences of strains isolated from water, but smaller than the ones originated from poultry meat (Table 1). We found the presence of a higher number of plasmid replicon sequences in *E. coli* strains originated from poultry meat (C4_38: 5 plasmid replicon sequences; C2_45: 3 plasmid replicon sequences) than in the strains obtained from water (E5895: 1 plasmid replicon sequence; E6003: no plasmid replicon sequence detected) or from *Daphnia obtusa* (ED1: 2 plasmid replicon sequences; ED157: no plasmid replicon sequence detected) (Supplementary file 1). Poultry meat strains had a higher number of virulence genes (C4_38: 32; C2_45: 29) than the *Daphnia* strains (ED1: 12; ED157: 16) and the water strains (E5895: 10; E6003: 12) (Supplementary file 1). Our analysis revealed that more phage sequences were present in ED1 and ED157 genomes than in the other genomes analysed here. Specifically, we found 7 phage sequences in ED1 and 5 in ED157 (Supplementary file 1), whereas only three and 0.5 phage sequences were found in poultry and in water strains, respectively.

**Figure 4.**
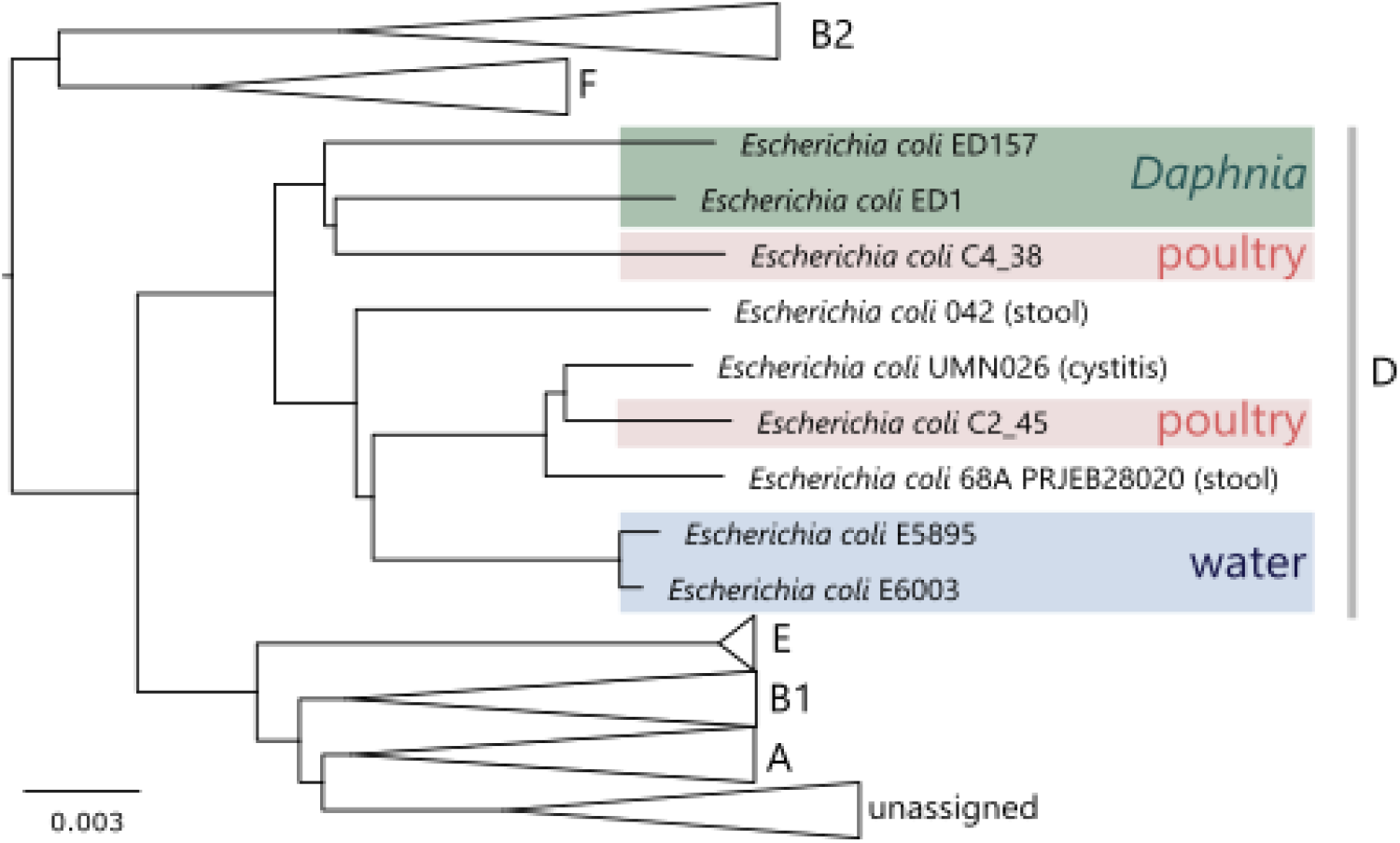
Phylogenetic tree of *E. coli* genomes included in tab. 1. The tree was constructed with MICROSCOPE tool with a neighbour-joining algorithm on Mash genomic distances. The scale bar represents 0.003 substitutions per nucleotide position.

To compare which genes were different in the *Daphnia* isolates compared to other *E. coli*, a pangenomic analyses was done with the strains listed in table 1 using the protein family sorter tool of PATRIC (Supplementary Table 3). The pangenome was composed of 7108 protein families while the core genome consisted of 3789 protein families (53,7%). *E. coli* isolated from poultry meat and from *Daphnia* shared 57.5% protein families (from a total of 6787 protein families), while *E. coli* isolated from *Daphnia* and from freshwater bodies shared 65% (from a total of 5993 protein families).

The genomes from all groups shared a very high number of protein families that can help the strains to thrive in the different habitats: we found, for instance, the presence of protein families related to the production of the capsular polysaccharides, or the presence of protein families linked to the Type I fimbriae system. Interestingly, we detected the presence of RhS protein families, which are supposed to inhibit the intracellular growth as primary function (Koskiniemi, et al. 2013) and the presence of some protein families linked to sucrose utilization only in isolates from *Daphnia* and poultry meat. Focusing on the Accessory genome of *E. coli* isolated from *Daphnia* (i.e. protein families that were not found in the other two groups), specific groups included e.g. the presence of xanthosine-related protein families, which allow bacteria to utilize purine nucleoside as carbon and energy source, and the presence of poly-beta-1,6-N-acetyl-D-glucosamine (PGA) protein families, which are involved in the synthesis, the export and the localization of PGA polymer, a necessary component for the formation of biofilms, protecting the bacteria to adverse environmental conditions.

### 3.4 *Interaction of* E. coli *with* Daphnia obtusa

#### Attachment

First we verified where *E. coli* was localised in the animal by incubating ED1-gfp and ED157-gfp strains separately with live *Daphnia for 4 hours*, dissecting the animals and performing a qPCR assay targeting the *gfp*-gene to verify the presence of *E. coli* on the various body parts (Figure 1A). We found that 70 ± 8% of the administered ED1-*gfp* and ED157-*gfp* were found in the gut compared to filter apparatus and carapax (data in Supplementary Table 4).

We then tested whether *E. coli* was digested by *Daphnia* or if *Daphnia* functioned as a refuge for *E. coli* or if they simply passed through the gut. We incubated ED1-gfp and ED157-gfp with *Daphnia* for 4 hours and then transferred them to new clean water (Figure 1B). Compared to the control (transferred water without *Daphnia*) we found a significantly higher abundance of both *E. coli* strains in the surrounding water of *Daphnia* (Figure 5).

**Figure 5.**
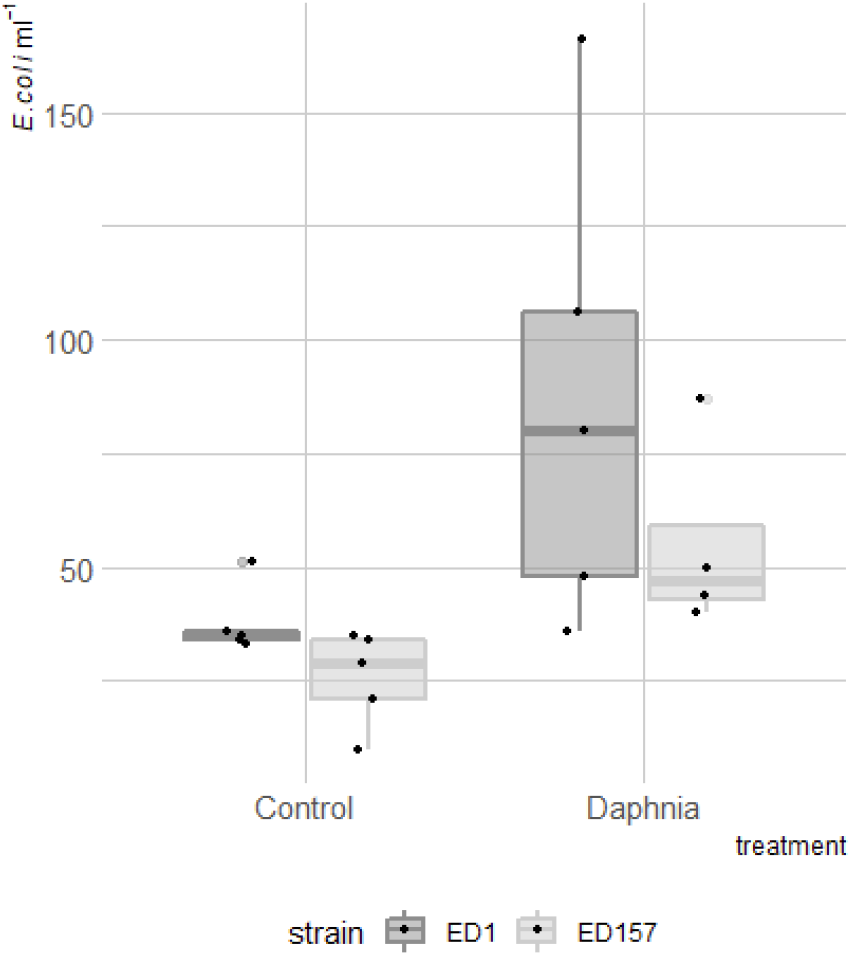
*E. coli* density in the treatment where only water was transferred (Control) and where animals were transferred (*Daphnia*) to sterile water after feeding on *E. coli* strains ED1-gfp (dark grey) or ED157-gfp (light grey). For each plot, the tick horizontal line represents the median value, the box includes 50% of the data from the first to the third quartile, the whiskers extend to the minimum and maximum data within the 1.5 interquartile range and the grey dots represent single outlier data points outside such range. The original data points for each treatment are superimposed on the plots as jittered black dots.

#### *Persistence with and without* Daphnia

We investigated how the presence of *Daphnia* impacted on the general survival of ED1-*gfp* in freshwater systems in continuous culture experiments (chemostat). We therefore filled eight chemostat vessels with ALW, a phytoplankton culture, *E. coli* and incubated *Daphnia* at different densities (0, 2, 4, 8, 10, 15 animals per vessel, Figure 1C). We monitored the abundance of culturable *E. coli* overtime in the water and found that the number of *Daphnia* had a slightly significant negative effect on the abundance of *E. coli* (glm: Estimate = −0.11±0.06, z = −2, p = 0.048). However, culturable *E. coli* abundances were in the same order of magnitude with only 1-6 CFU detected per ml of surrounding water in all treatments after 10 days of incubation even without animals (Figure 6). Some animals were then washed and guts dissected and *E. coli* cells were regrown in a plate reader to see whether there were culturable *E. coli* cells in the gut of the animals. One fourth of the total 12 dissected adult *Daphnia* (2 from vessel 5 and 1 from vessel 3, 0 from vessel 6 and 1) resulted in growth of *E. coli* within the first 48h of incubation, whereas no growth was detected from the gut of juvenile animals (total of 9 animals, Supplementary data 2).

**Figure 6.**
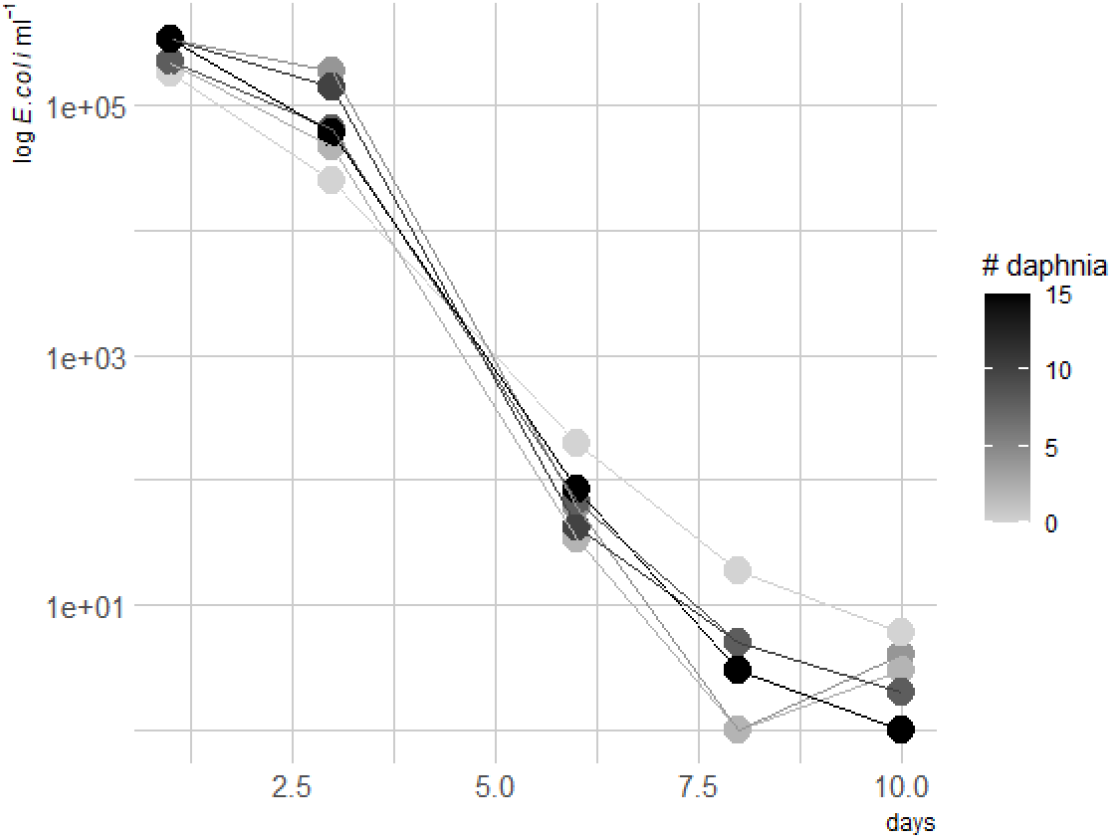
Log transformed cell density of tagged *E. coli* in chemostat experiment over time. The shade of grey of the dots and lines indicates the starting number of *Daphnia obtusa* added to the vessels.

#### Coexistence of ED1 and ED157

We then conducted an additional experiment where we combined both strains ED1-*gfp* with ED157-DsRed and ED157-*gfp* with ED157-DsRed, respectively, and incubated them either without animals, alive animals, dissected guts or dissected carapax (Figure 1D). At day 6 and 8 the abundances of culturable ED1 and ED157 (average of both tag combinations) were not statistically different (Figure 7, Table 2) whereas differences in treatments were visible: all treatments were different from each other except for the treatment with no *Daphnia* and the treatment with carapax pieces (Table 2). Living animals caused a faster reduction of *E. coli* abundances in the surrounding water than the other treatments. *E. coli* growth with carapax pieces increased in numbers notwithstanding the presence of other bacteria and reached numbers that were very similar to the treatment that contained only the tagged-strains. In the presence of gut pieces and naturally associated gut bacteria the abundances of both strains were reduced much more rapidly and at the end their abundance was similar to the one with live *Daphnia* (Figure 7).

**Figure 7.**
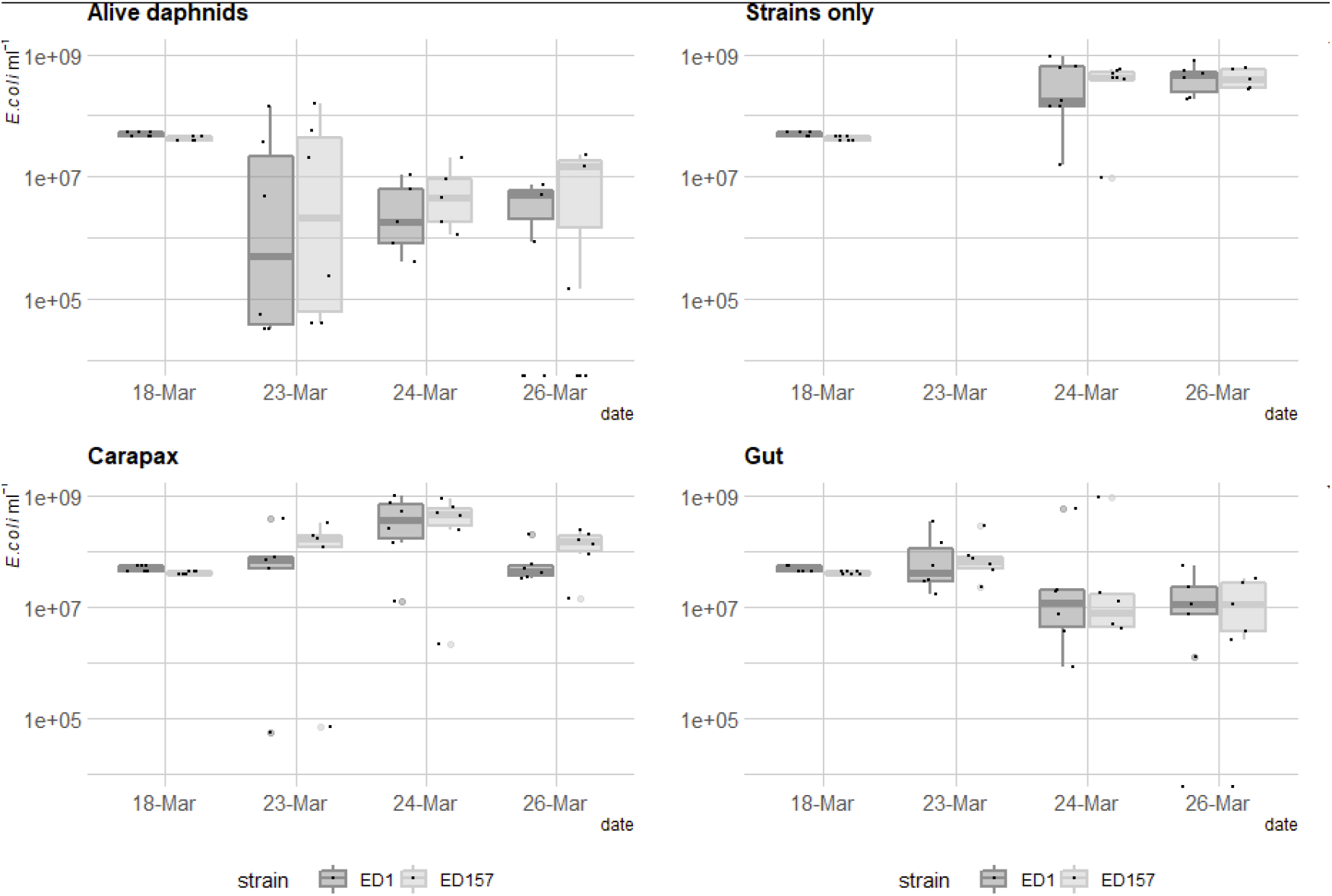
*E. coli* cell density over time in a batch experiment with the addition of *Daphnia obtusa* (Alive daphnids), no daphnids (strains only), pieces of *Daphnia* carapax (Carapax) and gut pieces (gut). Dark grey shows ED1 and light grey ED157 strains labelled with fluorescent proteins.

**Tab. 2.**
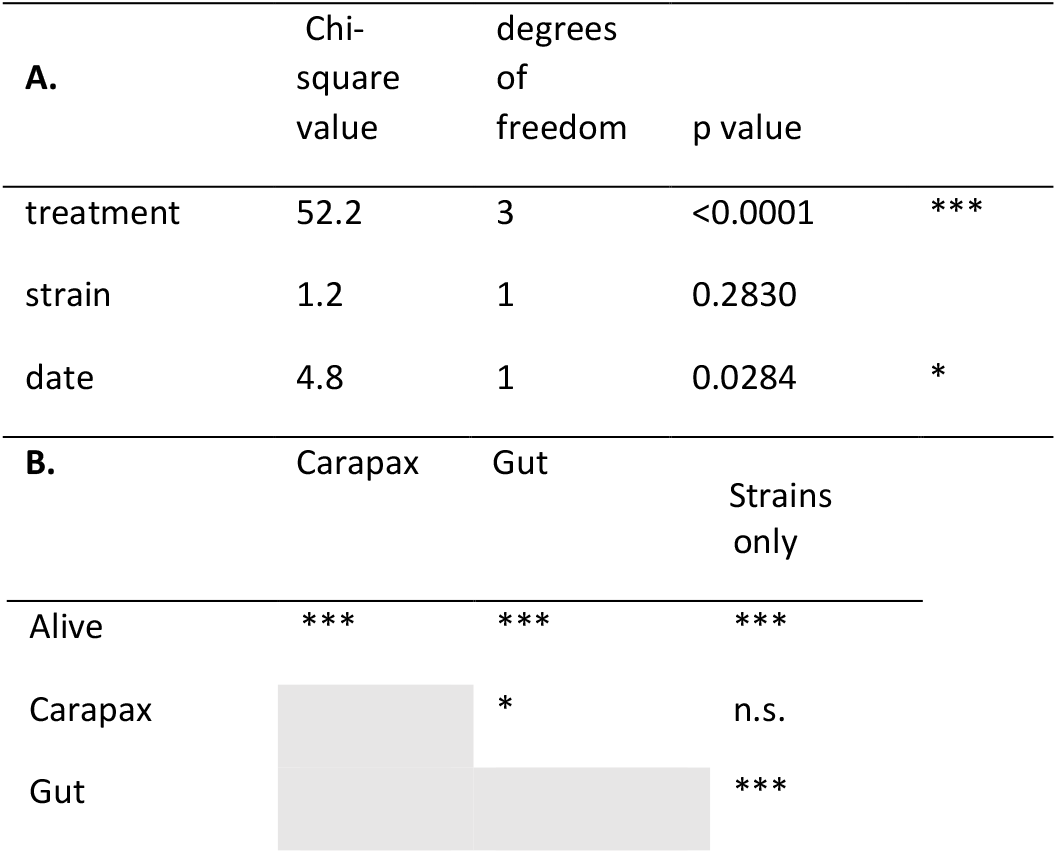
A. Statistical output from the generalised linear model made for the coexistence experiment to evaluate the dependence of the abundance of *E. coli* on the treatment (four levels: alive, carapax, gut, and strains only), the strain (two levels: ED1 and ED157) and the sampling date (two levels: May 24^th^ and 26^th^). The table is a type-II analysis-of-variance table with Wald chi-square tests for predictors. B. Significance of the differences in the pairwise comparisons between the four treatments from a Tukey post hoc test. *** means p value < 0.001, * p value < 0.05 and n.s. p value > 0.05.

## 4. Discussion

In this study we tackled the question whether *Daphnia* was a host for *E. coli* in freshwaters. While on one hand these bacteria are known to not be competitive in such environments and to be grazed easily when entering freshwaters through faecal pollution (i.e. (González, et al. 1992; Eckert, et al. 2019)), the evolution of freshwater *E. coli* (Touchon, et al. 2020) and renaturation of these bacteria have been observed (Ishii, Ksoll, et al. 2006; Ishii and Sadowsky 2008), showing that at least part of the incoming *E. coli* must survive over longer time periods in water. In a previous study we found that lake zooplankton could show remarkable quantities of *E. coli* related 16S rRNA gene sequences and we confirmed this here by quantifying the abundance of *uid*A gene, an unambiguous indicator for *E. coli* presence (Figure 2) (Eckert, et al. 2020). Indeed, this is not the first time that *E. coli* is found associated with zooplankton (Ishii, Yan, et al. 2006). Evidence is accumulating that freshwater zooplankton microbiota is rather flexible in terms of composition (Callens, et al. 2015; Macke, et al. 2020; Eckert, et al. 2021; Pfenning-Butterworth, et al. 2021), meaning that the association with zooplankton might be an interesting potential niche for the short-term survival of FIB. Such a habitat offers protection from protistan grazing and higher nutrient concentrations thanks to filtration feeding from the animal (Eckert and Pernthaler 2014). Furthermore, surface attachment is generally considered favourable for the survival of such bacteria compared to a planktonic lifestyle (Costerton, et al. 1999; Allen, et al. 2010). We thus tested here whether the association with *Daphnia* allowed *E. coli* to survive longer in aquatic habitats compared to when the animal was not present. In this study we found that *Daphnia* can function as a short-term and transitional host for *E. coli*: as hypothesised, *Daphnia* did reduce abundances of *E. coli* in the surrounding waters but it was not responsible for the complete removal of *E. coli*, since many bacterial cells survived gut passages (Figure 5) and *E. coli* was still detected after 10 days of co-culturing with *Daphnia* (Figures 5 and 6). In this study we confirm that *E. coli* localizes mainly in the gut of *Daphnia* and that at least part of its population survives the gut passage (Burnet, et al. 2017; Ismail, et al. 2019). However, the association with *Daphnia obtusa* did not seem to give a long term advantage in the survival of *E. coli* (Figures 6&7).

Many studies have recently suggested to use *Daphnia* as a biological controlling mechanism for *E. coli* contaminations in water: through experiments using very large densities of animals and bacteria these studies showed that the abundance of *E. coli* was reduced due to grazing of *Daphnia* (Nørgaard and Roslev 2016; Burnet, et al. 2017; Ismail, et al. 2019). The addition of *Daphnia* is surely feasible to reduce large abundances of *E. coli*, but here we showed that *E. coli* persisted even in the presence of *Daphnia* in low abundances that were more similar to those found in nature. In fact, we also showed that *E. coli* was still culturable from the gut, even if the bacterium was in very low abundance in the surrounding water. In our environmental survey we found it associated with different zooplankton hosts, especially cladocerans and rotifers (Figure 2), but particularly abundant in a specific sample of zooplankton from Lake Maggiore (Figure 2). However, also other samples from Lake Maggiore were analysed in the large study of zooplankton-associated microbes (Eckert, et al. 2021), and we did not find any *E. coli* in these. This shows that the short-term association can also occur in nature and that such association might consequently also spread *E. coli* that enter the system through superficial contamination due to the animals vertical and horizontal migration (Grossart, et al. 2010). However, there does not seem to be an actual persistence of these bacteria, indicating that the occurrences in the gut are rather stochastic events and that *E. coli* does not form part of the general *Daphnia* microbiota.

In the experiment where we incubated *E. coli* with dissected guts we observed a similar reduction of the bacteria (after 8 days) as was seen with alive *Daphnia* (Figure 7). This data strongly indicates that the competition with the gut microbiota was the main reason for reduced abundances of *E. coli*. Another interesting finding was that *E. coli* seemed to profit from the presence of carapax pieces of *Daphnia*, which are composed mostly of chitin. Both strains grew better in the presence of carapax, despite they did not have chitinolytic enzymes in their genome. It is more likely that the two strains indirectly profited from chitin degradation since such degradation is usually more efficient when done by multiple species (Corno, et al. 2015) and many bacteria are known to profit from these compounds without being directly involved in the primary degradation (Beier and Bertilsson 2013). The presence of poly-beta-1,6-N-acetyl-D-glucosamine (PGA) protein families, which are involved in the synthesis, the export and the localization of PGA polymer, shows that these *E. coli* strains might also be involved in multispecies biofilm formation (Kang, et al. 2018).

We isolated *E. coli* from *Daphnia* collected from a small pond, however with major difficulty. Despite multiple isolation campaigns and a clear presence of *E. coli*, confirmed by amplification of the *uid*A gene in these pond daphnids, we only isolated ten strains (Figure 3). This could mean that *E. coli* associated with *Daphnia* were in a viable but nonculturable state (VBNC), which has been observed in other freshwater environments (Liu, et al. 2008) or when exposed to sunlight (Pommepuy, et al. 1996). For *Enterococcus faecalis* it was shown that much higher numbers are detected attached to plankton with culture-independent methods, compared to the culturable fraction of these bacteria, and it has been suggested that this attachment in a VBNC state is a mode of survival of this species in freshwater (Signoretto, et al. 2004). A similar situation might also be true for *E. coli*.

The *E. coli* strains isolated from *Daphnia* in this study belonged to phylogroups D/E or B1. The analysis of our *Daphnia* deriving *E. coli* strains themselves did not give strong indication that these were environmental strains. Touchon and colleagues have shown that freshwater *E. coli* strains had a reduced genome (Touchon, et al. 2020), a typical form of adaptation to oligotrophic environments (Baumgartner, et al. 2017; Salcher, et al. 2019). In an experimental system Baumgartner and colleagues showed that such genome reduction was rather fast when bacteria were under predation (only a few hundred generations, Baumgartner, et al. 2017 (Baumgartner, et al. 2016)). In the case of our *E. coli* strains their genome was of intermediate size, meaning that they were smaller than the genomes found in poultry meat derived *E. coli*, but larger than those of freshwater *E. coli*, which could indicate a certain transition to adaptation of the genome. The two analysed *Daphnia*-associated *E. coli* genomes did not cluster with the freshwater isolates but with those isolated from poultry meat. In fact also the *E. coli* strains isolated from freshwater might derive from avian faeces (Meerburg, et al. 2011) and survive associated with zooplankton for a short time. The *E. coli* genomes also showed some traits that were considered important to adapt to different environments and for survival in freshwater, e.g., the presence of protein families related to the production of the capsular polysaccharides, which protect the bacteria from several environmental stress factors (Walk, et al. 2007; Azurmendi, et al. 2020), or the presence of protein families linked to the Type I fimbriae system, which allows bacteria to attach to several eukaryotic cells (Gally, et al. 1993). Whether or not zooplankton is a place of genetic adaptation of FIB to the environment is an interesting question arising from this study.

Overall our results showed that the FIB *E. coli*, when released into the aquatic environment, can form a short-term association with zooplankton, e.g. *Daphnia*. We demonstrated that *E. coli* does not belong to the core microbiota of *Daphnia*, suffers from competition by the natural microbiota of *Daphnia*, but may resist passages through its gut and profit of its carapax to survive in water. This association did not prolong their long-term survival in our experiments, but might provide a niche where these bacteria can encounter other aquatic bacteria, a possible spot for horizontal gene transfer, and a possible spot for genomic adaptation.

## Acknowledgments

PJC-Y was supported by a APOSTD/2019/009 Post-Doctoral Fellowship from Generalitat Valenciana. Research on Lake Maggiore was funded by the International Commission for the Protection of Italian-Swiss Waters (CIPAIS).

## Author contribution

EME, ADC, DF and GC designed the study. ADC and NC isolated *E. coli*, VM and CC characterised *E. coli* phenotypes, CV, GM, NC and BC characterised *E. coli* by MLST, PJCY, FR and EC analysed *E. coli* genomes, SB, EC and FM marked *E. coli* strains, EME and DF analysed 16S amplicon sequencing data, EME, DF, ADC, GB, WQ and GC conducted laboratory experiments, ADC and EE wrote the article with contribution from all the authors.

